# GEF Cytohesin-2/ARNO: a novel bridge between cell migration and immunoregulation in synovial fibroblasts

**DOI:** 10.1101/2021.10.04.463120

**Authors:** Yilin Wang, Çağlar Çil, Margaret M. Harnett, Miguel A. Pineda

## Abstract

The guanine nucleotide exchange factor cytohesin-2 (ARNO) is a major activator of the small GTPase ARF6, and has been shown to play an important role(s) in cell adhesion, migration and cytoskeleton reorganization in various cell types and models of disease. Interestingly, dysregulated cell migration, in tandem with hyper-inflammatory responses, is one of the hallmarks associated with activated synovial fibroblasts (SFs) during chronic inflammatory joint diseases, like rheumatoid arthritis. The role of ARNO in this process was unknown but we hypothesized that the pro-inflammatory milieu of inflamed joints induces local activation of ARNO-mediated pathways in SFs, promoting an invasive cell phenotype that ultimately leads to bone and cartilage damage. Thus, we used small interference RNA to investigate the impact of ARNO on the pathological migration and inflammatory responses of murine SFs, revealing a fully functional ARNO-ARF6 pathway in SFs, which can be rapidly activated by IL-1β. Such activation promotes cell migration and formation of focal adhesions. Unexpectedly, ARNO was also shown to modulate SF-inflammatory responses, dictating the precise cytokine and chemokine expression profile. Our results uncover a novel role for ARNO in SF-dependent inflammation, that potentially links pathogenic migration with initiation of local joint inflammation, offering new approaches for targeting the fibroblast compartment in chronic arthritis and joint disease.

## Introduction

Fibroblasts are stromal cells of mesenchymal origin that, in addition to their well-described structural roles, are essential for maintaining tissue homeostasis, as they orchestrate local immunity and inflammation, promote wound healing and control matrix remodeling (1). Thus, dysregulation of fibroblast activity by genetic, epigenetic or even environmental factors can prime pathological networks leading to chronic disease and autoimmunity (2). Such pathogenic fibroblast activation has been demonstrated in inflammatory Rheumatoid Arthritis (RA), a chronic inflammatory disease affecting the joints.

In healthy conditions, fibroblasts are essential to provide nutritional support and preserve joint homeostasis. Thus, fibroblasts cooperate with tissue-resident macrophages to form the synovium, a unique membranous organ lining the joint cavity whose barrier function maintains immune privilege in the joint (3). However, during RA, synovial fibroblasts (SFs) undergo a pathogenic transformation that destabilizes the normal function of the synovium and triggers inflammatory mechanisms, such as recruitment of immune cells, production of inflammatory cytokines (IL-6, CCL2, MCSF, RANKL), formation of ectopic lymphoid-like structures (4–6) and responses to inflammatory factors (IL-1β, TNF, IL-17). As a result, inflammatory SFs establish self-perpetuating inflammatory circuits based on secreted factors. Additionally, SFs-dependent pathogenesis is associated with changes in their adhesion and migration capacities, which ultimately creates the pannus, an overgrowth of the synovium observed only in arthritic joints that invades and damages bone and cartilage. As also observed in tumorogenesis, pathological SF migration is a result of dysregulated expression of matrix degrading enzymes, like MMPs, and aberrant upregulation of adhesion molecules, such as cadherin-11, CD82 and integrins or focal adhesion kinase (FAK) (7–9). The characterisation of functional heterogeneity provided by single-cell RNA-Sequencing (RNA-Seq) has allowed a rapid advance in the understanding of the pathophysiology of SFs in RA (10–12), but the mechanisms that underpin SFs migration and invasiveness are still unclear. This could be because Next Generation Sequencing (NGS) does not take into account the influence of the extracellular matrix and/or post-translational regulatory mechanisms governing cell signalling, the impact of which on cell function would not be detected by transcriptomic approaches alone.

Relating to these cell communication aspects, ADP-ribosylation factors (ARFs) are a family of small GTPases with roles in modulating membrane structure and the intracellular vesicle transport required for cell motility and attachment of cells to matrix components (13–16). ARF function is post-translationally regulated by guanine nucleotide exchange factors (GEFs), exchanging GDP for GTP to generate the active GTP-ARF. One particular GEF, cytohesin-2 (Cyth-2), also known as ARF nucleotide-binding site opener (ARNO), has been shown to promote cellular adhesion and migration in several cell types, including epithelial in cell migration, ARNO-dependent activation of ARF6 induces recycling of integrin β1 (17) and modulates actin remodeling (18,19) even in the absence of its GEF activity (20). Interestingly, Zhu et al. (21) also showed that ARNO acts as a branch point, separating IL-1β-dependent effects on cytokine secretion and barrier permeability in endothelial cells. Specifically, IL-1β triggers the MyD88-ARNO-ARF6 axis to compromise the integrity of the endothelial biological barrier, diverging from the canonical IL-1β-MyD88-IKK-NFkB pathway that promotes cytokine secretion and inflammation. Exposure of mice undergoing Collagen-Induced Arthritis (CIA) to the cytohesin inhibitor SecinH3 reduced disease severity and vascular permeability in the joints (21), but regulatory effects of ARNO on SFs in an inflammatory environment cannot be ruled out.

We thus hypothesized that inflammatory cytokines in the arthritic joint upregulate expression of ARNO in SFs, in turn activating ARF6-dependent pathways that promote the invasive phenotype characteristic of arthritic SFs. Thus, ARNO could offer a novel molecular switch discriminating between inflammatory and invasive fibroblasts in the arthritic synovium. Consistent with this hypothesis, we observed that ARNO was up-regulated upon IL-1β stimulation in SFs, ex vivo. Likewise, ARNO was found to be necessary to induce the morphological changes required for SFs migration. However, in contrast to the responses reported for endothelial cells (21) ARNO is also involved in regulation of SF inflammatory cytokine secretion. Collectively, our results indicate that the role of ARNO as a molecular checkpoint between inflammation and disruption of barrier permeability is not a universal function of this regulatory protein but rather depends on the cell type and anatomical location. Nevertheless, these data indicate that ARNO may provide a novel site for targeted intervention against joint disease in RA.

## Methods

### ex vivo culture of SFs

murine SFs were expanded *ex vivo* as described previously (22). Briefly, joint tissue was dissected from mouse paws and subjected to collagenase digestion for 80 minutes at 37°C. Cells were then cultured for 24 h to allow attachment to flasks and subsequently expanded in DMEM supplemented with 10% fetal calf serum (FCS,), 1% penicillin/streptomycin, 1% glutamine and 1% non-essential amino acid (all Invitrogen, UK) in 5% CO2 at 37 °C. Any contaminating myeloid cells were removed from cultures prior to experimental setup by negative selection using a biotinylated anti-CD11b antibody (Biolegend, #101204) followed by subsequent Streptavidin MicroBeads (Miltenyi Biotec, UK) for magnetic separation. Purity of SFs cultures (>99.5%) was assessed by positive expression of podoplanin by Flow Cytometry using anti-Podoplanin-Alexa Fluor 647 (#156204, Biolegend, UK) and anti-CD11b-FITC (#11-0112-85, Invitrogen, UK). For in vitro stimulation of SFs, recombinant IL-1β, TNFα and IL-17 (Immunotools, Germany) were used for the indicated times at 10 ng/ml. STAT3 activation was inhibited in vitro using Cpd188 (Sigma-Aldrich,UK) at 73 μM.

### Short interference RNA (siRNA)

ARNO siRNA (Mm_Pscd2_3) or a negative non-specific siRNA control (Allstar siRNA) were transfected into SFs using HiPerFect transfection reagent (all Qiagen,UK) according to the manufacturer’s protocol. Briefly, siRNAs were diluted in 8% HiPerFect Transfection Reagent in DMEM (Invitrogen, UK) and incubated for 10-15 minutes at room temperature to form transfection complexes before adding to the cells (final siRNA concentration 10 nM). Cells were incubated with transfection complex for 24 hours and then washed. Cells were transfected again using this protocol after 3 days, prior to experimental analysis as indicated.

### Collagen-Induced arthritis (CIA) mouse model

Mice (8-10 weeks old male DBA/1 mice) were obtained from Envigo (UK) and maintained in the Biological Service Facility at University of Glasgow in line with the guideline of Home Office UK and AWERB, the ethics review board of the University, and licenses PIL IF5AC4409, PPL P8C60C865. Mice were immunized with type II bovine collagen (MD Biosciences) emulsified in complete Freund’s adjuvant (CFA, day 0), and then challenged with type II bovine collagen (in PBS) on day 21. Body weight, clinical scores and paw thickness were measured every two days. Disease scores were measured in a scale from 0-4 for each paw. Animals reaching an overall score of 10 or more were immediately euthanized

### RNA isolation and qRT-PCR

RNA from SFs was extracted using either EZ-10 RNA Mini-Preps Kit (Bio Basic, USA) or RNeasy Micro Kit (Qiagen, UK) according to manufactures’ instructions. cDNA was synthesized with High-Capacity cDNA Reverse Transcription Kit (Thermo Fisher Scientific, UK). Gene expression was measured by RT-qPCR which was performed using either KiCqStart® SYBR® Green Primers (sigma-aldrich, UK) or TaqMan™ Gene Expression Assay (Thermofisher, UK) as indicated. Actin was used as an endogenous control to normalise samples and relative gene expression was evaluated using the comparative C_t_ (ΔΔC_t_) method. Sybr green primers used were actin/NM_007393, Cxcl9/NM_008599, Ccl9/NM_011338 and Cldn1/NM_016674 whilst Taqman mRNA primers were used for Actb (Mm02619580_g1), ARNO (Mm00441008_m1), IL-6 (Mm00446190_m1), CCL2 (Mm00441242_m1), MMP3 (Mm00440295_m1), MMP13 (Mm00439491_m1), TNFRSF11b (Mm00435454_m1) and TNFSF11 (Mm0041906_m1).

### Immunofluorescence staining

Cells seeded on chamber slide were incubated in Fixation buffer (Biolegend, UK) for 10 min at room temperature, washed three times with PBS, permeabilised (0.5% Triton X-100 in PBS) for 10 min and treated with blocking buffer (PBS 1% BSA or 10% normal serum from the species in which the secondary was raised) prior to incubation with primary antibodies overnight at 4 °C. The following primary antibodies were used: anti-vinculin (V9264, Sigma, UK), anti-CDH11 (71-7600, Invitrogen, UK), Alexa Fluor™ 488 Phalloidin (A12379, Invitrogen, UK). Samples were then incubated with donkey anti-mouse Alexa fluor 647 (A31571, Invitrogen, UK) or goat anti-rabbit Alexa fluor 647 (A21245, Invitrogen, UK) for 1h at RT. Slides were mounted with SlowFade™ Diamond Antifade Mountant with DAPI (S36968, Invitrogen, UK) to counterstain nuclei. Samples were visualized using either EVOS (EVOS™ FL Auto 2, Thermofisher, UK) or Zeiss LSM 880 confocal microscopes. Magnification?

### ARF6-GTP pull down assay

Activation of ARF6 (ARF6-GTP) was quantitated using the Arf6 Pull-Down Activation Assay Biochem Kit from Cytoskeleton (BK033-S, USA) according to manufacturer’s instructions. Briefly, cells were lysed on ice in 200 μl Cell Lysis buffer and centrifuged at 10,000 × g for 1 min at 4°C. An aliquot (20 μg) of lysate was saved for Western blot quantitation of total ARF6 expression. Then, cell lysate (125 μg) was added to 5 μg GGA3-PBD beads (Cytoskeleton, USA) to pull down ARFs-GTP and incubated at 4°C on a rotator for 1h and then the beads were extensively washed to remove unbound protein. The beads were then resuspended and boiled in Laemmli buffer (SigmaAldrich, UK) and the samples resolved by SDS-PAGE prior to detection of ARF6 by Western blot.

### Western blotting

Cells were harvested and lysed in RIPA buffer (Thermo Fisher Scientific, UK) containing protease inhibitor and PhosSTOP tablets (Roche, UK) and sample protein concentration determined using the Pierce® BCA Protein Assay Kit (Thermo scientific, UK). SDS-PAGE was then performed, and proteins were transferred onto nitrocellulose membranes. Membranes were blocked with TBS-T 5% non-fat milk prior to overnight incubation with for primary antibodies at 4 °C (used at 1:5000 unless stated otherwise). The following antibodies were used: anti-Cytohesin2 (sc-374640; Santa Cruz Biotechnology, 1:1000), anti-ARF6 (ARF-06; Cytoskeleton, 1:500), anti-p44/42 MAPK (CST #4695, UK), anti-GAPDH (CST #2118, UK), anti-P38 (CST #9212, UK), anti-P38 phospho (CST #9211, UK), anti-STAT3 (CST #9139, UK) and anti-STAT3 phospho (CST #9145, UK). Membranes were then incubated with anti-rabbit (CST #7074P2, UK) or anti-mouse (CST #7076s, UK) HRP-conjugated secondary antibodies for 1h at room temperature and washed three times in TBS-T. Expression signal was detected by ECL Western Blotting Substrate (Thermo scientific, UK) and the protein bands were quantified using GelAnalyzer 2010a software with the relative integrated density values normalised to ERK1/2 and GAPDH expression values.

### Cytokine detection by ELISA

Cell supernatants were collected for evaluation of secreted cytokines. SFs (10^4^) were seeded in 200 μl of complete DMEM in 96-well plates pre-coated with bovine fibronectin (R&D, UK). For siRNA silencing experiments, siRNA transfection was performed in 12-well plates, with the cells detached and re-seeded in 96-well plates when the siRNA transfection protocol was completed. Cells were cultured in the absence or presence of 10 ng/ml IL-1*β* for 24 hours, then supernatants were collected and frozen. ELISA kits (Dou set, R&D) were used to evaluate the levels of IL-6, MMP3 and CCL2 in the supernatants according to the manufacture’s instructions. Cell viability was checked using a MTS proliferation kit (abcam, UK).

### Cell migration assay

4-well μ-Dishes (#80466, Ibidi, UK) coated with fibronectin were seeded with 10,000 SFs. Cells were grown until confluency, and inserts were then removed. The space without cells was then measured as a reference for subsequent cell migration, and it was measured again after 24 hours to determine the distances migrated as calculated using ImageJ software.

### Flow cytometry

The following antibodies were used with flow cytometry: anti-Podoplanin-Alexa Flour 647 (#156204, Biolegend, UK), anti-CD11b-FITC (#11-0112-85, Invitrogen, UK), DAPI (#32670, Sigma, UK) and proliferation dye eFluor 670 (#65-0840-90, eBioscience, UK). Cells were trypsinized, stained at 4C and analysed using a LSR II flow cytometer (BD Biosciences). For proliferation studies, synovial fibroblasts were labelled with 10μM proliferation dye for 10min on ice. Labeling was stopped by adding 5 volumes of 10% FCS culture medium and incubate 5 minutes on ice. DAPI was used to discriminate live and dead cells. Data were analyzed with FlowJo software 10.7.1.

### RNAseq and bioinformatic analysis

Total RNA from cultured synovial fibroblasts was isolated using RNeasy Micro kit (Qiagen, Germany). RNA integrity was checked using the Agilent 2100 Bioanalyzer System, and the RIN value was >9 for all samples. Library preparation was done using the TruSeq mRNA stranded library preparation method and samples were sequenced (2 × 75 bp) with an average of 30 million reads. All RNA-seq reads were then aligned to the mouse reference genome (GRCM38) using Hisat2 version 2.1.0, and read counts were generated using Featurecounts version 1.4.6. Data quality control, non-expressed gene filtering, median ratio normalization (MRN) implemented in DESeq2 package, and identification of differentially expressed (DE) genes (padj < 0.01, |foldChange|>2) was done using the R Bioconductor project DEbrowser (23). Gene Ontology (GO) enrichment and KEGG pathway enrichment was conducted with Metascape (24).

### Statistical analysis

All statistical analysis was performed in GraphPad Prism 8. Data are presented as the mean ± standard error (SEM) with students t-test used to show differences between two study groups for unpaired samples. P values <0.05 were considered significant.

## Results & Discussion

### IL-1β activates the ARNO-ARF6 pathway in murine SFs

Synovial fibroblasts (SFs) were expanded from joint tissue collected from healthy animals as described previously (22) and their phenotype confirmed by their expression of podoplanin and the absence of the myeloid marker, CD11b (Figure 1A). To test our working hypothesis, we first evaluated expression of ARNO mRNA in SFs, both under resting conditions and upon stimulation with recombinant versions of pro-inflammatory cytokines typically found in the inflamed joint (Figure 1B). IL-1β, IL-17 and TNF were selected as they have been shown to be drivers of disease both in animal models (25,26) and in human RA, and have been associated with distinct pathotypes in RA (27). However, whilst all three cytokines upregulated IL-6 mRNA levels, only IL-1β significantly increased ARNO mRNA levels (Figure 1B) suggesting, in line with previous observations in endothelial cells (21), that IL-1β provides a specific signal(s) that promotes ARNO activity.

**Figure 1.**
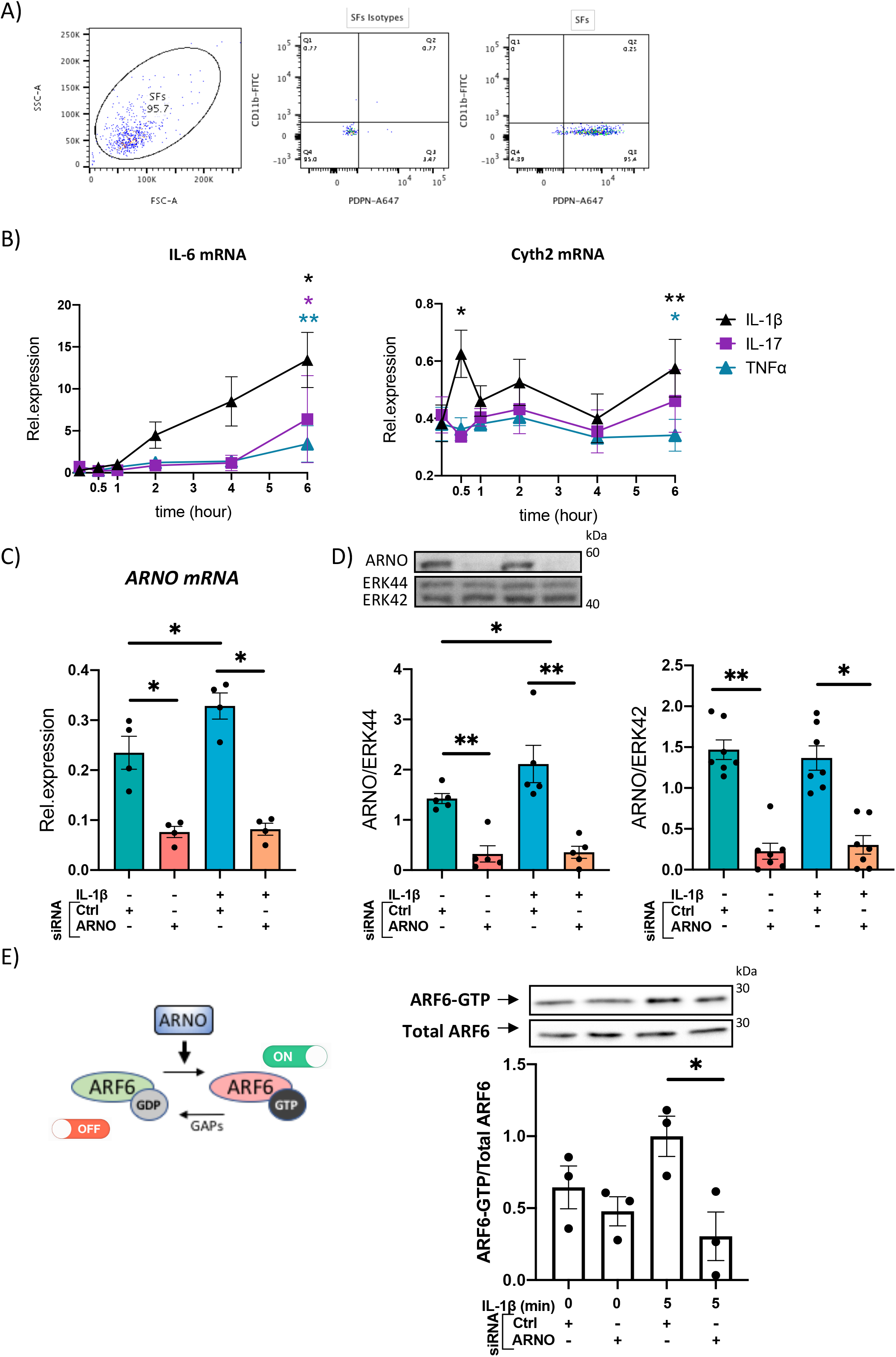
IL-1β upregulates ARNO in SFs. A) SFs were isolated from mouse synovium and expanded ex vivo. Cells were trypsinased and expression of podoplanin (PDPN) and CD11b was evaluated by flow cytometry. B) SFs were stimulated with IL-1β, IL-17 and TNFα (10 ng/ml) and RNA was extracted at the indicated time points (0, 0.5, 1, 2, 4 and 6 hours). Expression of ARNO and Il6 mRNA levels were quantified by RT-qPCR. C) Naïve SFs were transfected with ARNO or control Allstars siRNA, followed by stimulation with recombinant IL-1β (10 ng/ml) when indicated. RNA was extracted after 6 hours and ARNO mRNA expression was evaluated by RT-qPCR. D) SFs were transfected and stimulated as in (C), and proteins were extracted 24h after stimulation to conduct western blot. ARNO protein was detected by specific monoclonal antibodies, total brain lysate was loaded as positive control (data not shown) and total Erk was used as loading control. Image shows one representative experiment, and column graph shows quantification of band intensity for ARNO normalised with Erk. Each dots represents one independent experiment. E) SFs treated with Allstar or ARNO siRNA followed by IL-1β stimulation were assayed by ARF6-GTP pull down assay and subsequently immunoblotted with anti-ARF6 antibody. Relative activation of ARF6 was normalised to unstimulated control siRNA-treated samples. Each dot represents an independent experiment n≥3, error bars represent SEM, *p<0.05, **p<0.01 by Wilcoxon’s matched-paired signed rank test in B, by Mann-Whitney test in D and E.

To address the functional role of ARNO in SFs, we used small interfering RNA (siRNA) approaches which were highly effective at downregulating expression of both ARNO mRNA (78.3%±0.08), Figure 1C) and ARNO protein (79.1%±0.22, Figure 1D), relative to the negative control AllStars siRNA (Qiagen, siRNA with no homology to any known mammalian gene) in resting and IL-1β-stimulated SFs. ARNO activates ARF6 to compromise vascular permeability in arthritis and other inflammatory conditions (21,28,29) and it also triggers key effector mechanisms involved in cell migration (29–32). Thus, to investigate whether the ARNO-ARF6 axis was also active in SFs, we knocked-down ARNO expression by siRNA and evaluated activation of ARF6 (GTP-bound) in response to IL-1β (Figure 1E). This showed that ARNO is necessary for IL-1β to induce ARF6 activation, confirming the presence of the IL-1β-ARNO-ARF6 axis in SFs.

### ARNO is necessary for SF migration and focal adhesion formation

Next, we sought to characterise the physiological function(s) of ARNO in SFs. ARNO-ARF6 strongly causes cells to migrate in culture, through well-described mechanisms involving Rac1, DOCK180/Elmo and R-Ras and cytoskeleton-dependent changes in cell morphology (30,33). Increased ARNO activity also modifies the way that cells interact and move through the extracellular matrix, by controlling β1-integrin recycling (33,34). Because migration and integrin-dependent signaling are known hallmarks of inflammatory SFs, we evaluated the ability of SFs to migrate using wound healing assays (Figure 2A). SFs migration was significantly reduced when ARNO was knocked-down, in agreement with the function of ARNO in other cell types. We evaluated the effect on cell proliferation, but ARNO inhibition did not have a significant impact (Figure 2B), suggesting that distinct molecular mechanisms are likely responsible for dysregulation of cell migration and proliferation in RA. To initiate migration, SFs in RA must disassemble their cell contacts in the organized synovium and establish new ones with the extracellular matrix. The reduced migration of siARNO SFs led us to hypothesize that ARNO was involved in the formation of focal adhesions (FAs), multiprotein complexes that concentrate actin polymerization to control migration through engagement with the extracellular matrix. To investigate this, we stained SFs for vinculin, a protein that anchors actin to the membrane in FAs. We found that ARNO siRNA significantly reduced both the number and structure (length and area) of focal adhesions (Figure 2C). Overall, these data highlight the role that ARNO plays in determining the fundamental biology of SFs, including their morphology, cell adhesion and migration.

**Figure 2.**
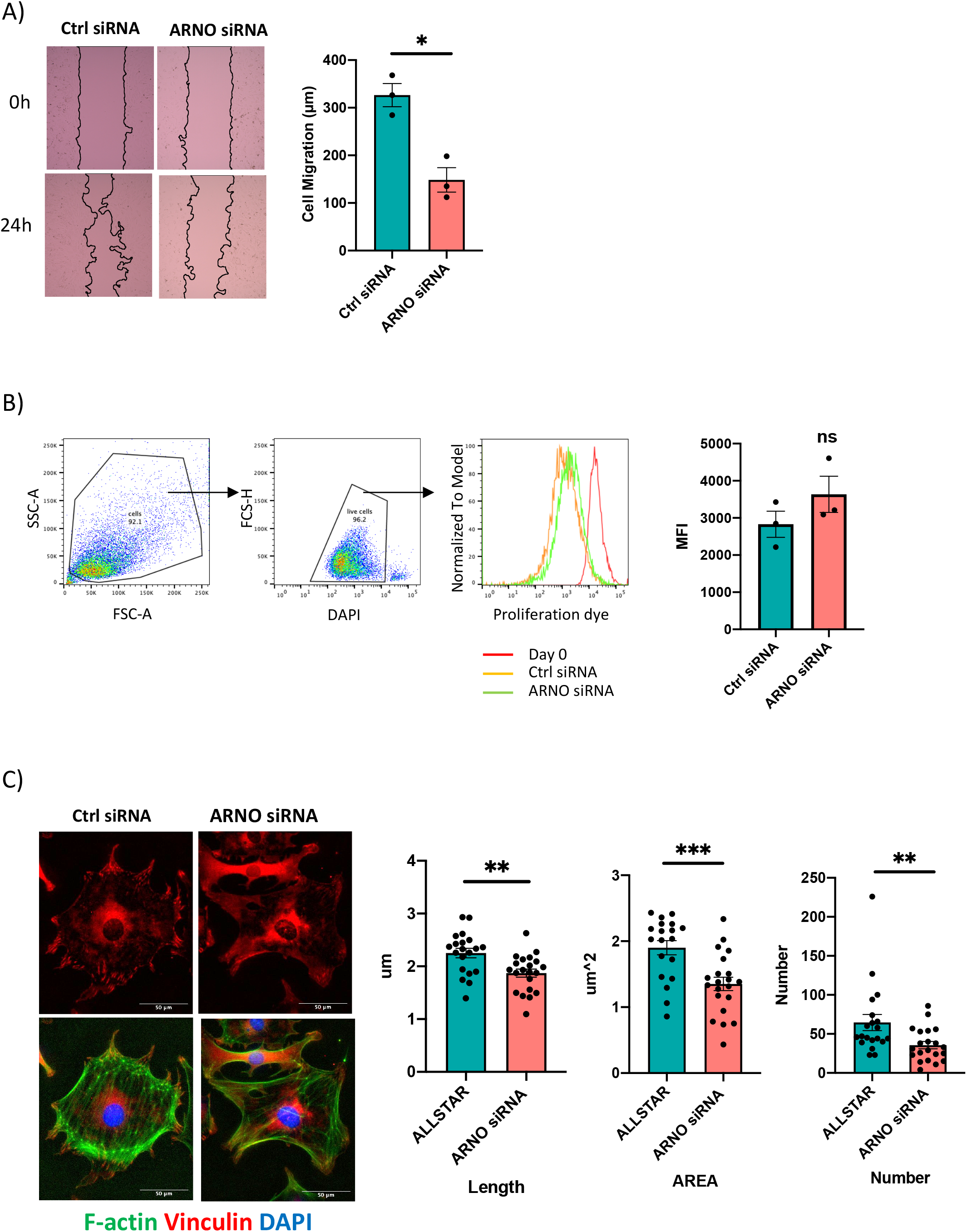
ARNO regulates SFs migration and assembly of focal adhesions. A) SFs were seeded in migration chambers and grown until monolayer confluence. Cells were then treated with either Allstars control or ARNO siRNAs, and inserts were removed to perform migration assays. Pictures show one representative experiment, superimposed black lines delineate the cell-free area. Bar chart shows the mean of cell migration distance ± SEM (n=3) calculated with ImageJ software. B) SFs were labelled with proliferation dye then analysed by flow cytometry or treated with control or ARNO siRNA and continued in culture. After 5 days, mean fluorescence intensity was evaluated in live cells (identified by dimmer DAPI staining). Histograms show one representative experiment, and mean fluorescence intensity (MFI) of three independent experiment is shown in the column graph where error bars represent ± SEM (n=3). C) Immunofluorescence staining of Vinculin (red) and F-actin (green) in SFs treated with Allstars control or ARNO siRNA. Scale bar: 50um. Vinculin length, area and number were analysed. Each dot represents a calculated value in a cell, error bars represent ± SEM (n=1). ns: not significant, *p<0.05, **p<0.01, ***p<0.001 by Mann-Whitney test.

### ARNO is required for cytokine secretion in activated SFs

In RA, initiation of SF migratory programs likely directly links with inflammation because interactions with the matrix through up-regulated cell-to-cell adhesion molecules can increase secretion of cytokines and matrix metalloproteinases (MMPs) (35–37). Thus, although ARNO is not often reported as an inflammatory mediator, we proposed that formation of ARNO-dependent complexes upon IL-1β stimulation triggers secretion of inflammatory cytokines, coupling pathological invasiveness with local SF-derived inflammation. Consistent with this, we observed that reduction in ARNO expression (using siRNA) substantially inhibited IL-1β-dependent IL-6 and CCL2 secretion by naïve SFs (Figure 3A) and down-regulation of MMP3 was also observed (Figure 3A), albeit this did not reach statistical significance. To assess the prospect of using ARNO as a therapeutic target, it was important to establish whether the responses observed with naïve SFs also pertained in the chronically activated cells generated during disease. Thus, SFs explant cultures from animals undergoing Collagen-Induced Arthritis (CIA) were examined for ARNO-dependent expression of IL-6, CCL2 and MMP3 (Figure 3). As expected from previous studies (38), CIA SFs showed elevated cytokine production compared to naïve cells, corroborating their inflammatory phenotype. ARNO knock-down in such CIA cells reduced the IL-1β-dependent release IL-6, CCL2 and MMP3, the latter also in a significant manner in these cells. Collectively therefore, we observed that suppression of ARNO limited the ability of both naïve and CIA SFs to respond to IL-1β, indicating that ARNO-signalling may provide a novel link between inflammation and matrix remodeling/invasiveness in SFs.

**Figure 3:**
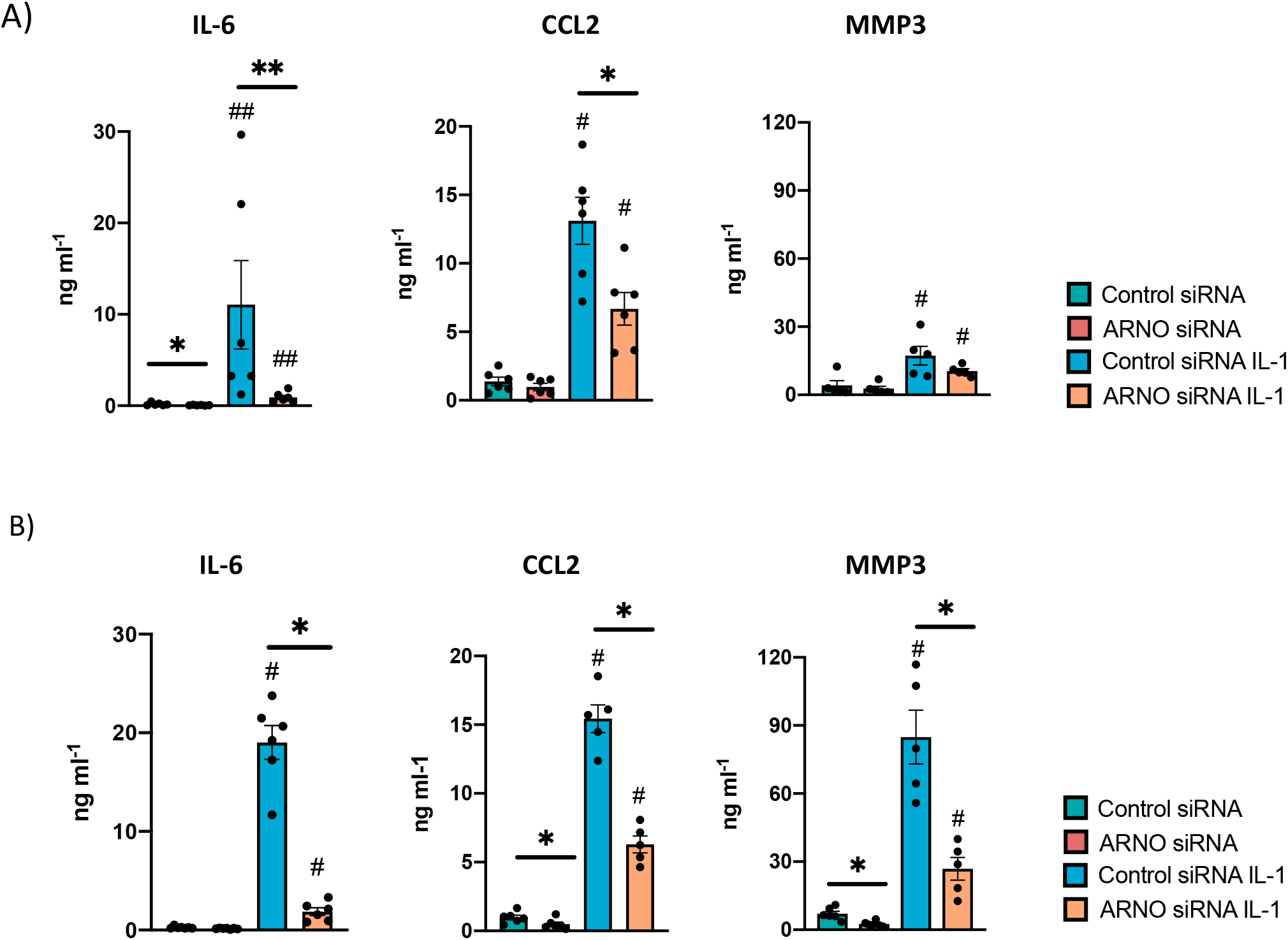
Reduction of ARNO expression down-regulates IL-1β-mediated cytokine expression. Secretion of IL-6, CCL2 and MMP3 was analysed by ELISA in the supernatant of control and ARNO siRNA naïve SFs (A) and SFs expanded from mice undergoing experimental Collagen-Induced Arthritis (B). Cells were stimulated with IL-1β as indicated, and cell supernatants were collected after 24h for cytokine quantification. Each dot represents one independent experiment analysed in triplicate, error bars represent SEM. *p<0.05, **p<0.01 versus respective siRNA control; #p<0.05 versus unstimulated control, Statistical significance was evaluated by Mann-Whitney test.

To obtain further mechanistic insight, we investigated potential signaling pathways regulated by ARNO (Figure 4). Thus, we explored the role of ARNO in the IL-1β STAT3 signalling pathway, as this can play key roles in both acute and chronic inflammatory responses upon IL-1β stimulation in SFs (38). We also evaluated the p38 MAPK pathway, as it has been shown to be important for RA pathogenesis and IL-1 signalling (39) and also for ARNO signaling in relation to MMP expression (31). Reduction in ARNO expression by siRNA transfection resulted in inhibition of STAT3 activation (as evidenced by its phosphorylation quantitated as pSTAT3/STAT3 expression) upon IL-1β stimulation, both in naïve (Figure 4A) and CIA SFs (Figure 4B). Of note, this inhibition did not reflect a downregulation of STAT3 expression. By contrast, no effect was observed on p38 activation or expression, ruling out a direct role for this kinase in ARNO-mediated inflammation and for ARNO in p38 coupling. To provide further support for a functional role for the IL-1-ARNO-STAT3 axis in SFs pathogenesis, we treated SFs with the specific STAT3 inhibitor Cpd188 (40), which efficiently blocked IL-1β-dependent activation of STAT3 (Figure 4C). Cpd188 treatment did not affect SF proliferation (Figure 4D), but it reduced migration in wound repair assays (Figure 4E), findings resembling the effect observed with ARNO knock-down (Figure 2A). Finally, we measured the effect of STAT3 inhibition on cytokine secretion upon IL-1β stimulation (Figure 4F). In this case, results did not fully recapitulate those observed in ARNO knock-down experiments (Figure 3). Thus, although STAT3 inhibition down-regulated CCL2 release, the levels of IL-6 and MMP3 remained unaltered (Figure 4F), findings suggesting that additional ARNO-dependent signals contribute to the production of these mediators.

**Figure 4:**
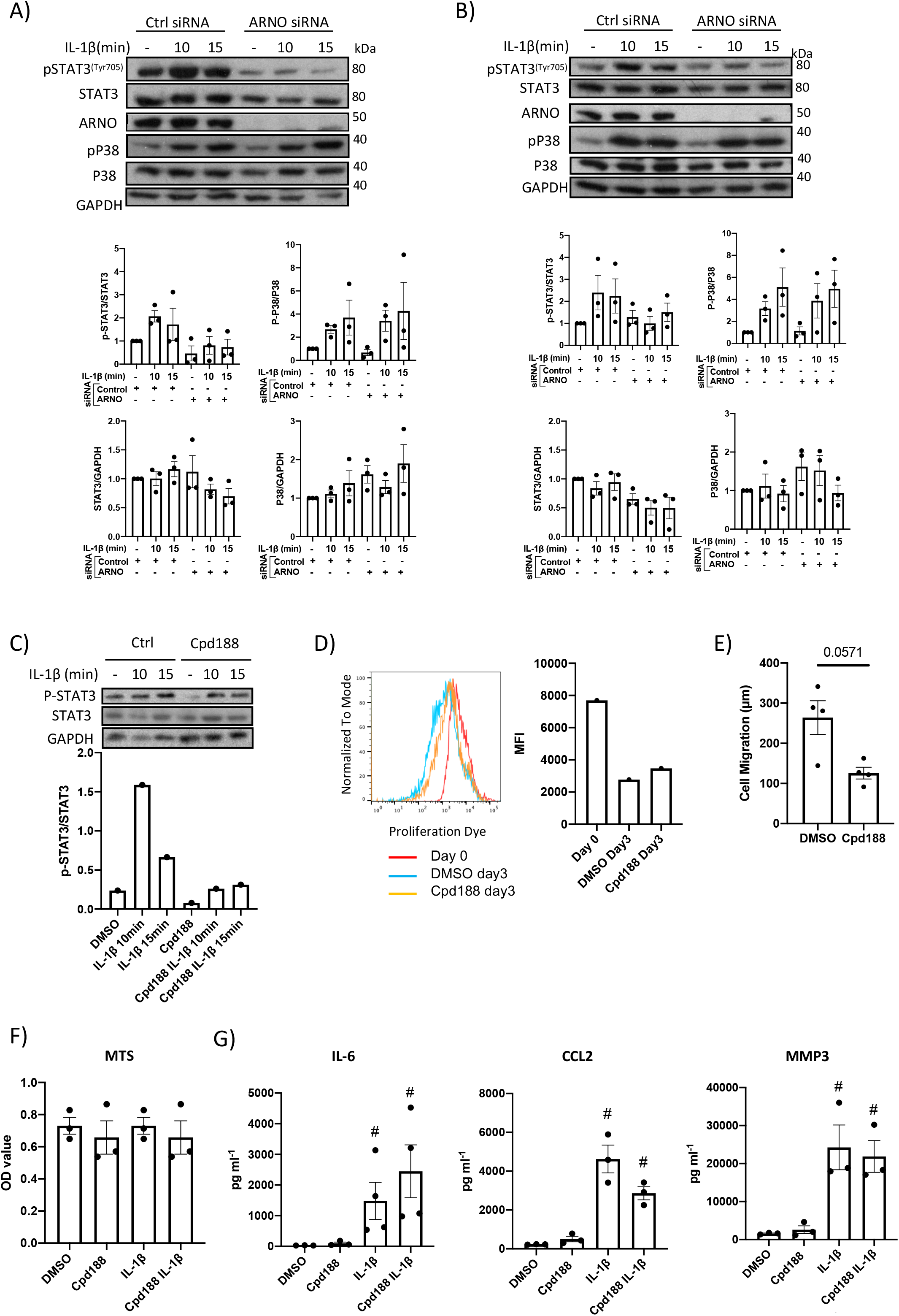
ARNO is required for STAT3 phosphorylation in SFs. Representative western blots of the naïve (A) and CIA (B) SFs treated with control and ARNO siRNA followed by IL-1β stimulation at indicated time. Anti-pSTAT3^Tyr705^, STAT3, ARNO, p-P38, P38 and GAPDH (sample control) antibodies were used. Quantification of phosphorylated/total and total STAT3 and P38 in naïve SFs and CIA SFs are presented as signal intensity normalised to GAPDH, error bars represent SEM (n=3). C) Representative western blots and quantification of phosphorylated and total STAT3 in control (DMSO) and Cpd188 (STAT3 inhibitor, 73uM) treated naïve SFs followed by IL-1β stimulation at indicated time (n=1). D) Mean fluorescence intensity of proliferation dye in SFs cultured for 3 days after treated with Cpd188 compared to control SFs. E) Migration distance of SFs was measured 24 hours after Cpd188 treatment, error bars represent SEM (n=4). Control or Cpd188 treated SFs followed by IL-1β stimulation as indicated for 24h, cell viability was assessed using MTS cell proliferation kit (F), IL-6, CCL2 and MMP3 in the supernatant was analysed by ELISA (G) error bars represent SEM (n=3). #p<0.05 versus unstimulated control, statistical significance was evaluated by Mann-Whitney test.

### ARNO-signaling fine-tunes SF-dependent inflammation upon IL-1β stimulation

To address identifying ARNO-associated pathways and hence further understand ARNO function in SFs, naive and IL-1β-stimulated cells, the latter treated with and without ARNO siRNA, were subjected to RNAseq analysis. Principal Component Analysis identified that the three groups displayed distinct transcriptome profiles (Figure 5A) and so we therefore compared the significant differential gene expression [foldchange > 4, adjp <0.01] existing amongst the three experimental groups (Figure 5B) to identify distinct transcriptomic signatures associated with each experimental group.

**Figure 5:**
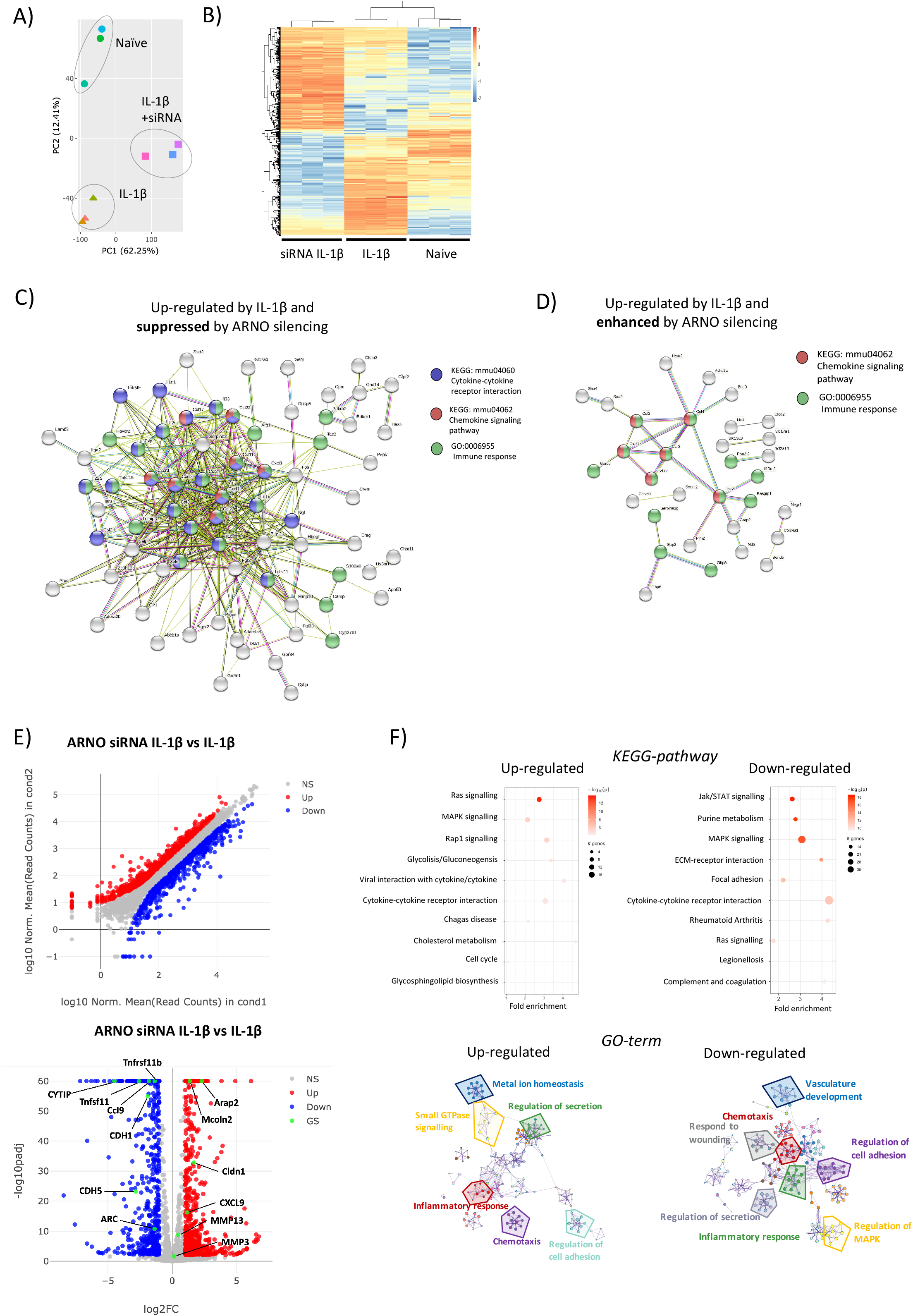
ARNO knocked-down SFs show a distinct transcriptomic profile in response to IL-1β stimulation. A) RNA was isolated (RIN>9) from naïve, IL-1β stimulated and IL-1β stimulated ARNO knocked down SFs (6 hours, n=3) and subjected to bulk RNA-Seq (75bp paired-end, 30M reads). Principal component analysis (PCA) is shown. B) Differential expression (DE) of genes amongst the three experimental groups. Genes were considered significant if they passed a threshold of padj < 0.01 and |log2foldChange| > 2 among any paired comparison. DE genes were then subjected to unsupervised hierarchical clustering and represented as Z scores. C,D) Function enrichment and network analysis regulated by IL-1β and ARNO expression. STRING protein-protein interaction network (https://string-db.org) was performed on DE genes from B, with genes up-regulated only in the IL-1β-treated group (C) or genes up-regulated only in the ARNO siRNA group (D). Significantly modulated pathways components associated with ARNO silencing upon IL-1β are shown. E) DE of genes in IL-1β stimulated SFs compared to ARNO siRNA IL-1β SFs. Genes are plotted as a scatter plot where x= gene expression in IL-1β SFs, y=gene expression in siRNA ARNO IL-1β SFs. Vulcano plot shows gene expression versus p value for each gene. Genes that pass a threshold of padj < 0.01 and |log2foldChange| > 2 in DE analysis are colored in blue when they are down regulated and red when they are upregulated in IL-1β treated cells. F) DE genes identified in (E) were used to conduct KEGG pathway enrichment analysis and GO-term pathway analysis. KEGG pathways are represented by circles and are plotted according to fold enrichment on the x-axis and −log10 p-value on the coloured scale. The size is proportional to the number of DE genes. Metascape enrichment network diagrams illustrate GO-term pathways significantly enriched for top up-regulated and down-regulated genes.

Firstly, we applied K-means clustering to our DE genes to identify which genes were up-regulated in IL-1β-stimulated, but not in ARNO siRNA IL-1β stimulated cells, to determine, which IL-1β-induced genes were suppressed by ARNO siRNA (Figure 5C). A list of 122 genes was generated, of which 72 (Supp. Table 1) were found to be functionally connected when subjected to String Protein-Protein Interaction Networks Functional Enrichment Analysis (41) (Figure 5C): the two most over-represented pathways were 1) CXC chemokines from the IL-8 superfamily, responsible for neutrophil and monocyte recruitment and 2) pro-inflammatory cytokines with crucial roles in joint inflammation, such as IL-6, CSF2, CSF3, IL-23 or TNFSF11 (RANKL). Interestingly, we found a significantly enriched number of cytokines induced downstream of JAK/STAT signalling (CSF2, CSF3, IL-6, IL-23a, LIF, Tslp), in line with the observed STAT3 activation by ARNO (Figure 4). Next, we similarly used the K-means approach to search for IL-1β-induced genes whose expression was enhanced when ARNO expression was silenced (Figure 5D) and this identifed 94 genes. Interestingly, this group included some pro-inflammatory chemokines (CCR3, CXCL13, CCL12, CCL3 and CCL4), although no inflammatory cytokines were found to be up-regulated by ARNO knock-down.

Overall, these results indicated that ARNO modulates IL-1β-driven inflammatory response rather than being essential for these functions. To examine the functional implications of such modulation, we directly compared IL-1β-stimulated SFs with those treated with IL-1β and ARNO siRNA SFs (Figure 5E), finding 384 up-regulated and 434 down-regulated genes [DE foldchange > 2, adjp <0.01] (supplementary table 2). In addition to the distinct cytokine and chemokine signature, this analysis revealed a significant modulation of genes involved in cell migration and cytoskeleton organization, such as Arc or cadherins (Cdh1, Cdh5) (42–44) as well as genes that can activate ARFs (Arap2, Cytip and Mcoln2) (45–47). Intriguingly, ARNO silencing also modified the expression of genes (Cmah, ST3Gal5, ST3Gal6, ST6Gal1) involved in the biosynthesis of sialylated glycoproteins, a pathway that we have recently shown to control SF activation in inflammatory arthritis (22). This DE gene list was investigated for pathway enrichment using Go-term and KEGG databases (Figure 5F) and, although signalling pathways are generally regulated by post-translational modifications, our RNAseq data suggest that MAPK and RAS signaling pathways were affected by ARNO knock-down, with up- and down-regulation of pathway elements being indicated, perhaps explaining the observed differences in cytokine receptor pathways. Critically, focal adhesion and JAK-STAT pathways were specifically down-regulated in ARNO siRNA SFs, in agreement with our functional results, whilst GO-term analysis also confirmed the role of ARNO in cell adhesion and migration in these cells.

### Dual role of ARNO in the regulation of pathogenic responses to IL-1β

Our transcriptomic data suggest that ARNO is involved in regulating SF responses to IL-1β, being essential for IL-6 production and for directing responses towards specific chemokines. To confirm and validate these results using RT-PCR. To further explore and confirm this, we selected a representative set of genes associated with inflammatory joint disease (refs), whose expression was either significantly suppressed or enhanced by ARNO knock-down in the RNASeq dataset: specifically, we selected 4 down-regulated genes, IL-6, Ccl9, Tnfsf11 and Tnfsf11b, and 4 up-regulated genes, MMP3, MMP13, CLDN1 and Cxcl9 (figure 6A) and validated their expression status by qRT-PCR. Whilst the cytokines, chemokines and MMPs were chosen for their pro-inflammatory and matrix remodelling roles, CLDN1 was selected as a tight junction transmembrane protein involved in cell-cell adhesion (48). In addition, TNFSF11 (TNF Superfamily Member 11 also known as Receptor activator of nuclear factor kappa-Β ligand, RANKL) is the major osteoclastogenic factor and TNFRSF11B (TNF Receptor Superfamily Member 11b, or Osteoprotegerin, OPG) acts as decoy receptor for RANKL and thus these genes are critical to the regulation of bone damage (49). Expression of these genes, plus ARNO (Cyth2) as control for ARNO silencing, was evaluated by qRT-PCR analysis of naïve murine SFs transfected with ARNO siRNA (or negative control Allstars siRNA) upon IL-1β stimulation as before. Although, as with the RNAseq experiment (Figure 6A), CLDN1, CXCL9 and Mmp13 expression was found to be up-regulated, levels of Mmp3 were unchanged in ARNO knocked-down cells after IL-1β stimulation (Figure 6B). Likewise, whilst levels of IL-6 and Ccl9 mRNA induced upon IL-1β activation were also reduced when ARNO was silenced, we did not observe any significant change in the expression of RANKL and OPG mRNA. However, as regulation of the RANKL/OPG axis by SFs has been shown to depend on the combined effects of inflammatory cytokines (50), we repeated the experiment using fibroblasts from arthritic CIA mice, which have become epigenetically modified in vivo as a result of their chronic exposure to multiple cytokines and proinflammatory mediators (51). Reflecting their imprinted pathogenic status, such CIA SFs showed a stronger ARNO up-regulation than naïve cells in response to IL-1β, but ARNO siRNA transfection was equally effective reducing ARNO mRNA expression (85.9%±0.05) (Figure 7A). Interestingly, RANKL induction after IL-1β stimulation was substantially reduced whilst that of OPG tended to be upregulated when ARNO expression was knocked-down in these CIA SFs and the differential regulation of these genes was highlighted by analysis of the ratio of their counter-regulatory expression (Figure 7B). Intriguingly, these data suggest that ARNO-dependent regulation of gene expression differs in Naïve and CIA SFs, with CIA SFs apparently more susceptible to adopting a pathogenic phenotype in response to the joint inflammatory mediator, IL-1β. Providing further support for this idea, whilst ARNO siRNA treated CIA SFs show a significantly reduced ability to produce IL-6 in both Naïve and CIA SFs, and Ccl2 ARNO is also shown to promote both MMP3 and MMP13 expression in CIA SFs (Figure 7C), whereas ARNO silencing had no effect on MMP3, and enhanced MMP13 synthesis in Naïve SFs.

**Figure 6:**
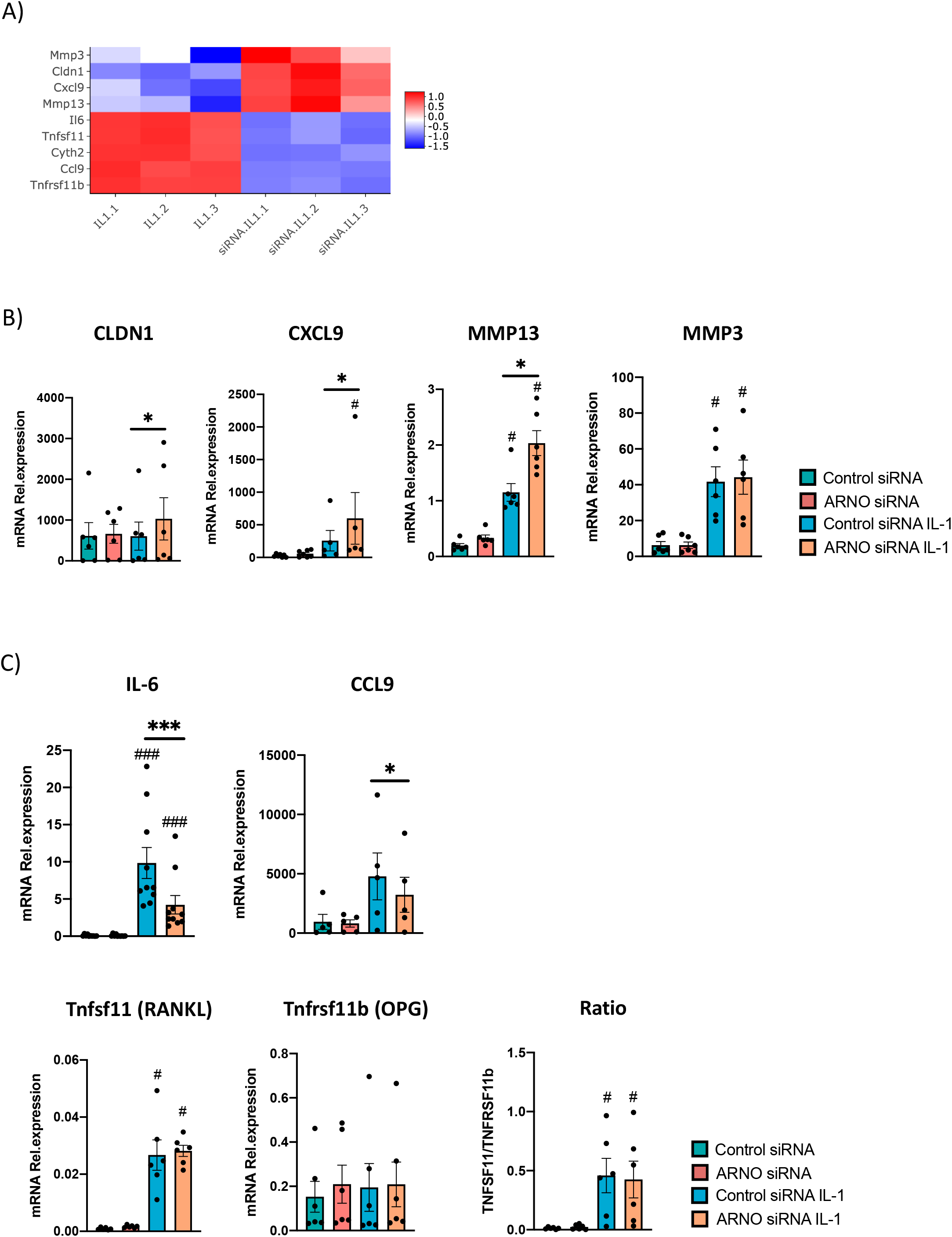
ARNO knock-down remodels the type of IL-1β-mediated inflammatory response. A) mRNA expression as detected by RNA-Seq from figure 5 for ARNO (Cyth2), IL-6, CXCL9, CCL9, CLDN1, MMP3, MMP13, TNFRSF11b and TNFSF11. B-C) RNA was isolated from IL-1β-stimulated and IL-1β-stimulated naïve SFs upon ARNO knock down by siRNA. Relative expression of genes shown in (A) was evaluated by RT-qPCR using the ΔΔC_t_ method and actin as housekeeping gene. B) ARNO, CLDN1, Cxcl9, MMP13, MMP3 mRNA relative expression. C) IL-6, Ccl9, TNFRSF11b and TNFSF11 mRNA relative expression. For B-C, each dot represents one independent experiment analysed in triplicate, error bars represent SEM (n≥5), *p<0.05, **p<0.01, statistical significance was evaluated by Mann-Whitney test.

**Figure 7.**
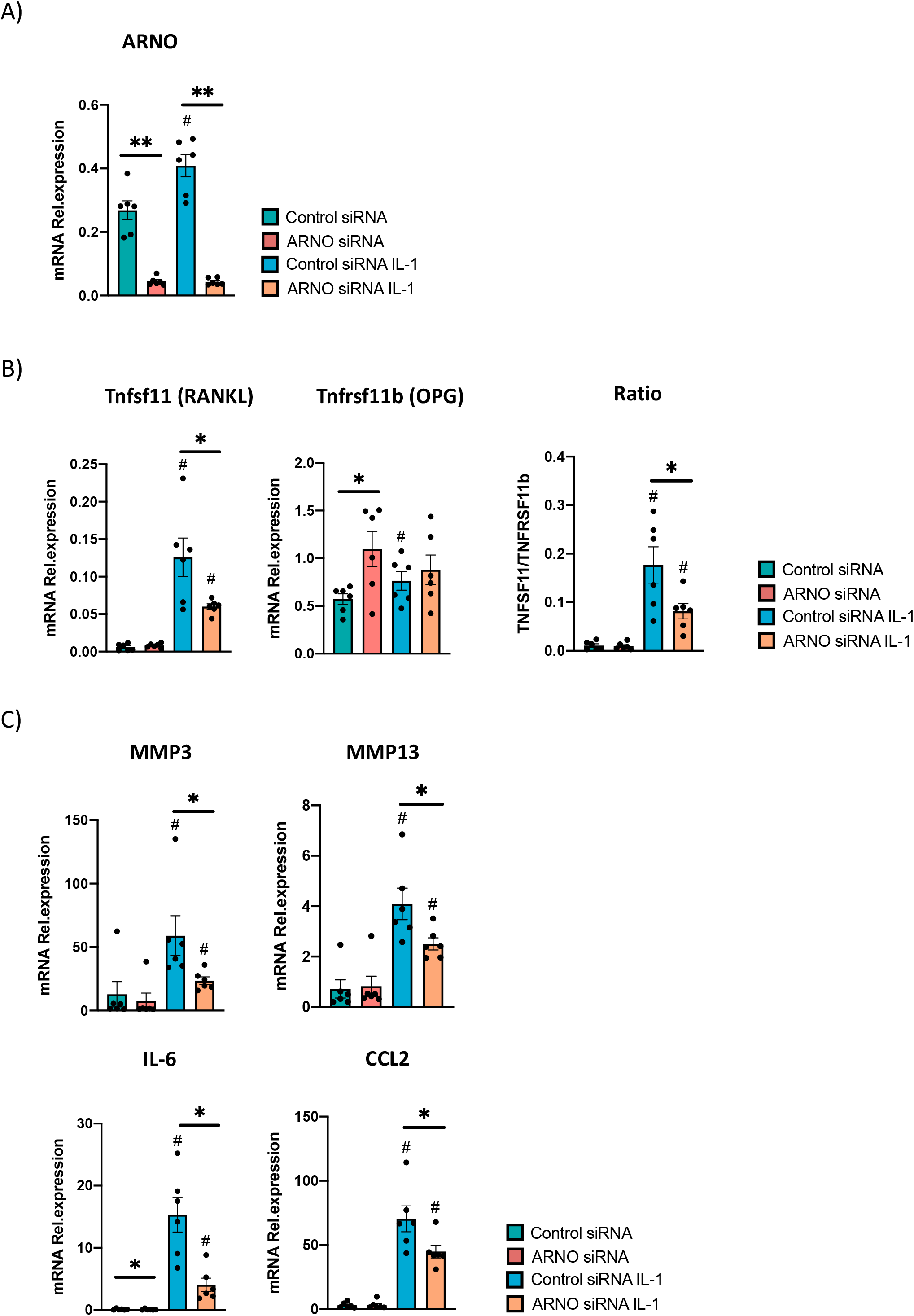
ARNO knock-down decreases IL-1β-dependent responses of CIA SFs. SFs were expanded from mice undergoing Collagen-Induced Arthritis. RNA was isolated from IL-1β-stimulated and IL-1β-stimulated (6 hours) SFs upon ARNO knock-down by siRNA. ARNO (A) TNFRSF11b and TNFSF11 (B) and MMP3, MMP13, IL-6 and Ccl2 relative mRNA expression was evaluated by RT-qPCR using the ΔΔC_t_ method and actin as housekeeping gene. Each dot represents one independent experiment analysed in triplicate, error bars represent SEM (n=6), *p<0.05, **p<0.01, statistical significance was evaluated by Mann-Whitney test.

Overall, we report here that i) ARNO is necessary for migration and reorganisation of focal adhesions in synovial fibroblasts and ii) a novel role of ARNO in modulating the inflammatory response of SFs in response to IL-1β. This represents a direct connection between the phathophisiological migration of inflammatory fibroblasts with their ability to initiate and modulate local immune responses. Interestingly, ARNO did not act as a switch for inflammation, but it modulated the type of inflammatory signals released by SFs. This role for ARNO in this specific cell type may offer new alternatives to modulate synovial fibroblasts-mediated immunity in chronic inflammatory joint disease, such as RA, without severely compromise systemic immune responses.

## Acknowledgements

The work was funded by a Career Development award to M.A.P. from Versus Arthritis (21221) and a CSC Scholarship awarded to Y.W.

---

**Table S1:** Up-regulated by IL-1β and suppressed by ARNO silencing

| Up-regulated by IL-1β and suppressed by ARNO silencing<br>(ARNO siRNA IL-1β Vs IL-1β) |  |  |  |  |  |
| --- | --- | --- | --- | --- | --- |
| ID | log2FoldChange | padj | ID | log2FoldCh | padj |
| Has1 | -4.737604979 | 9.92E-49 | Fos | -2.1771158 | 2.36E-298 |
| Ptgs2 | -4.555676183 | 1.05E-176 | Havcr2 | -2.1395481 | 4.46E-171 |
| Cytip | -4.496149816 | 0 | S100a8 | -2.0887185 | 6.19E-18 |
| Olr1 | -4.243993103 | 6.11E-163 | Ptger2 | -2.0667794 | 1.21E-44 |
| Gem | -3.948463743 | 1.44E-267 | Cxcl3 | -2.0501893 | 1.80E-159 |
| Lif | -3.941519452 | 0 | Csf2rb | -2.0267885 | 0 |
| Il23a | -3.755006442 | 2.56E-21 | Ier3 | -2.0262747 | 9.08E-181 |
| Tnfsf15 | -3.404873016 | 1.20E-221 | Fgf23 | -1.9825951 | 9.33E-20 |
| Arg1 | -3.376618335 | 8.28E-48 | Bdkrb1 | -1.7227577 | 1.06E-69 |
| Ereg | -3.30930589 | 0 | Mmp10 | -1.6636681 | 2.27E-43 |
| Sstr2 | -3.181799437 | 9.71E-25 | Cxcl2 | -1.6634727 | 9.47E-91 |
| Il1rl1 | -3.065438924 | 0 | Ngf | -1.5748574 | 5.11E-82 |
| Bdkrb2 | -2.97789197 | 1.86E-126 | Tslp | -1.5357421 | 2.89E-31 |
| Il1b | -2.928576937 | 1.48E-213 | Csf3 | -1.4771581 | 1.87E-38 |
| Abcb1a | -2.868626541 | 9.92E-117 | Cyp27b1 | -1.3870835 | 0.00033173 |
| Il1a | -2.863214444 | 7.14E-17 | Sele | -1.3176288 | 0.00475707 |
| Il33 | -2.81653226 | 0 | Lamb3 | -1.3078873 | 1.63E-08 |
| Dusp5 | -2.734788527 | 6.95E-214 | Tnfrsf9 | -1.2037322 | 0.00119434 |
| Procr | -2.673649006 | 6.06E-248 | Selp | -1.1932895 | 1.48E-07 |
| Fgf2 | -2.646084722 | 4.49E-210 | Ptges | -1.1429392 | 2.82E-66 |
| Hbegf | -2.63029002 | 1.75E-194 | Tac1 | -1.0944233 | 6.22E-06 |
| Cpm | -2.609931439 | 4.88E-43 | Adora2b | -1.0660125 | 1.17E-49 |
| Tnfsf11 | -2.583168423 | 3.67E-119 | Ccl11 | -1.0007754 | 6.27E-09 |
| Serpinb2 | -2.561595012 | 4.57E-275 | Gpr84 | -0.9867981 | 1.60E-45 |
| Gfpt2 | -2.554135392 | 0 | Adamts4 | -0.9234278 | 3.53E-31 |
| Ccl22 | -2.466230507 | 0.00049672 | Dkk1 | -0.6408036 | 0.38210511 |
| Grem1 | -2.449697728 | 9.66E-211 | Cxcl1 | -0.5978533 | 3.33E-20 |
| Hs3st1 | -2.447256142 | 4.22E-48 | Camp | -0.5811888 | 0.00126357 |
| Il6 | -2.399603961 | 1.10E-174 | Ccl17 | -0.5708553 | 0.09994657 |
| Gna14 | -2.3731284 | 0.00055922 | Il2ra | -0.5612924 | 0.2845768 |
| Apold1 | -2.359692273 | 6.46E-08 | Csf2 | -0.4474356 | 0.06856092 |
| Clstn3 | -2.331866004 | 7.34E-16 | Zc3h12a | -0.3870827 | 3.93E-11 |
| Chst11 | -2.322910993 | 0 | Ccl2 | -0.3246935 | 1.22E-08 |
| Crem | -2.303403204 | 1.31E-167 | Slc7a2 | -0.22676 | 0.00247768 |
| Itga2 | -2.230034594 | 4.36E-13 | Tnfaip3 | -0.1420346 | 0.04474146 |
| Penk | -2.196216083 | 1.91E-216 | Cxcr3 | -0.1041643 | 0.70532254 |

**Table S2:**
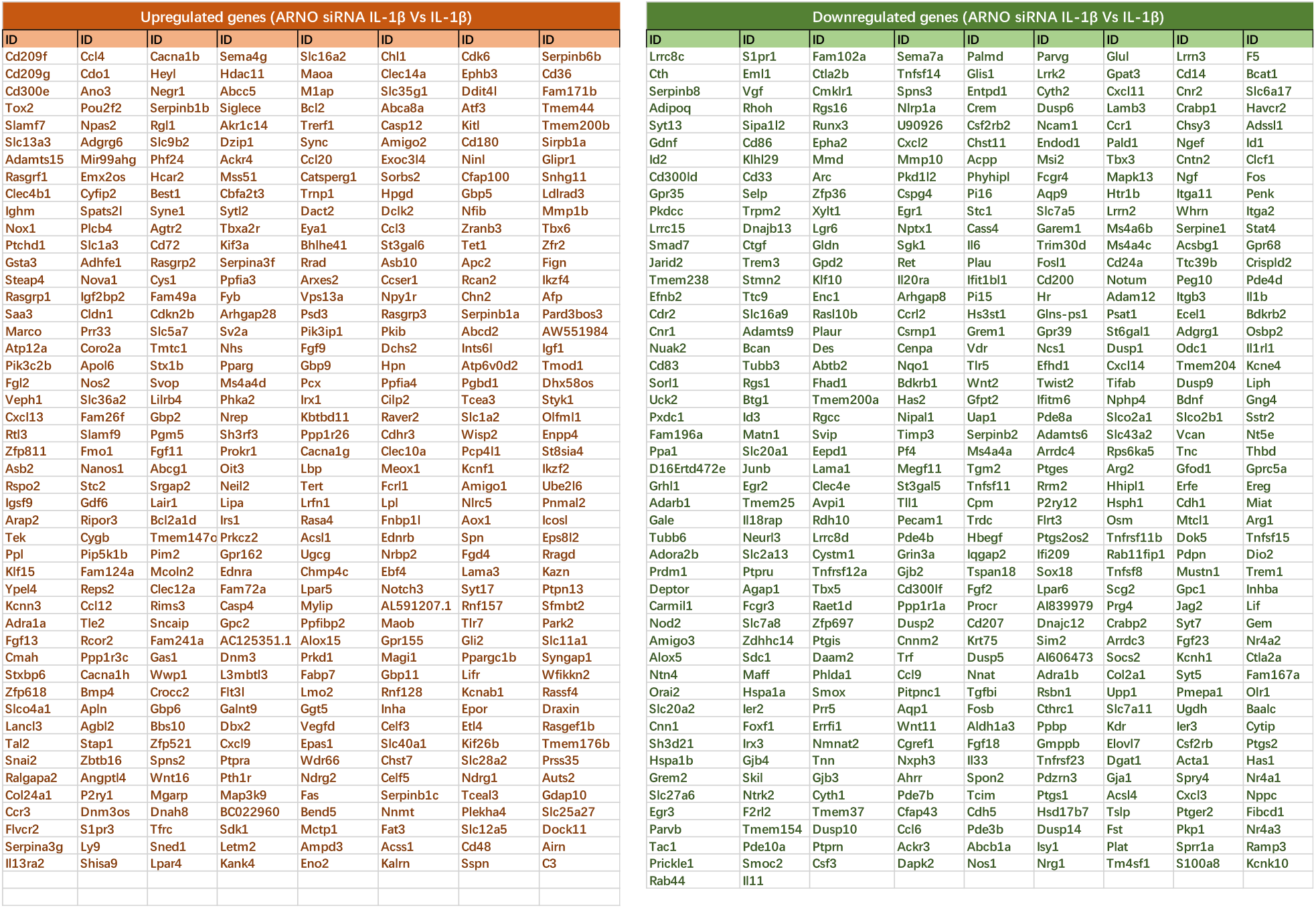
Up (left) and down (right) regulated genes compared 1β with 1β and ARNO siRNA SFs

